# *Alopecosa nagpag* acts on cardiac ventricular myocytes to kill prey

**DOI:** 10.1101/2023.03.27.534449

**Authors:** Zhixin Gu, Chenbo Long, Yuehua Lu, Biao Huang

## Abstract

Spiders are excellent predator to kill their prey by peptide toxins from its venoms. *Alopecosa nagpag* (*A. nagpag*) is a new identified wolf spider distributing in Yunnan province and nothing has known about the venom. In this study, venom of *A. nagpag* showed mild toxicity to Kunming mouse with LD_50_ of 3.32 mg/kg. Action potential duration (APD) was prolonged in a frequency-dependent manner and whole currents of neonatal rat ventricular myocytes (NRVMs) were inhibited by venom. Meanwhile, venom of *A. nagpag* could largely increase L calcium currents (I_CaL_). Whereas sodium current (I_Na_) and rapidly activating delayed rectifier potassium current (I_Kr_) were significantly decreased by 100 μg/mL venoms. No obvious inhibition was found on other ion channels such as rapidly activating and inactivating transient inward (I_K1_), rapid (I_Kr_) and slow (I_Ks_). As those ion channels play critical role in rhythm of cardiac ventricular myocytes, *A. nagpag* may lead prey to death by changing cardiac rhythm.

## 1. Introduction

Various ion channels on membrane of cardiomyocyte are the key factors on the generation and spreading of cardiac action potential (AP), which regulates cardiac rhythm and normal function of heart [1]. Due to the distinct contributions of different ion channel, ionic fluxes could generate action potentials in excitable cells[2]. There are six main ion channels expressed in cardiac ventricular myocytes: transient outward K^+^ current (I_to1_), delayed rectifier K^+^ current (I_Kr_ and I_Ks_) and inward rectifier K^+^ currents (I_K1_), voltage-gated Na^+^ current (I_Na_) and L-type Ca^2+^ currents (I_CaL_) [3].I_CaL_ inhibition can shorten the action potential and underlying I_CaL_ help to maintain the plateau phase of ventricular action potential. Slowing the inactivation of Na^+^ or Ca^2+^ currents or accelerating the inactivation of K^+^ current may contribute to prolong the action potential and to generate heart diseases, such as torsade de pointes or LQT [4]. Na^+^ channel blockers, procainamide prolongs ventricular repolarization and has been developed as class Ia antiarrhythmic drug. Lidocaine shortens action potential duration and is thought to be class Ib antiarrhythmic drug [5].

Spiders contains an extreme diversity of peptide neurotoxins acting on ion channel especially on sodium, potassium and calcium channels [6]. Wolf spiders are wide distribution in both coastal and inland. *Alopecosa nagpag* (*A. nagpag*) is a new wolf spider species distributing in Yunnan province. It lives in small cave without spin webs and hunt alone. When pouncing upon prey, it injects venoms to prey with chelicerae. Symptoms of its venomous bite include swelling, mild pain, and itching. It has supposed that *A. nagpag* contains large amounts of toxin, which was used to kill preys. However, nothing was known about the venom of *A. nagpag*. Here, we investigated the actvity of this venom on neonatal rat ventricular myocytes (NRVMs).

## 2. Materials and methods

### 2.1. Collection of the venom

Adult A. nagpag were kept in plastic box with plastic net and given water daily. An electro-pulse stimulator with a 25–50 V and 15–40 Hz pulse current was used to collected crude venom by stimulation of the chelicerae [19]. Crude venom was lyophilized and then stored in -80 °C until using.

### 2.2. Toxicity Assays

Venoms were dissolved in saline. Kunming mouse (30) were divided into five groups and each mouse was injected with 100 μL venom (0.5, 1, 2.5, 5 and 10 mg/kg). The value of LD50 (i.e. the dose lethal to 50% of animals) was fit with dose-response data by the following equation: y=(a-b/1+(x/LD50)n)+b, where x is the toxin concentration, y represents the percentage of deaths in the sample population, n is the variable slope factor, a is maximum response, and b is minimum response [20]. The animal study was reviewed and approved by the Animal Care and Use Committee at Kunming Institute of Zoology, Chinese Academy of Sciences (SMKX-20210516).

### 2.3. Cell isolation

Ventricular of two days neonatal rats Neonatal rat was used to dissociat ventricular myocytes (NRVMs) cell [21]. Firstly, it was cut into pieces on ice. Then, ventricular was continue digested with collagenase (Sigma) and trypsin (Sigma). The NRVMs cells were cultured in Dulbecco’s modified Eagle’s medium (DMEM)/F-12 culture medium with 10% fetal bovine serum [22]. The NRVMs cell pellets were incubated at 37 °C in a 95% O2 incubator for 1.5 h to separate non-cardiac myocytes. The NRVMs were continue cultured for 1-2 d for ion current recordings.

### 2.4. Electrophysiological recording

Whole-cell patch-clamp recordings were performed to record currents of various ion channels at room temperature[23]. For recordings, The capillary pipettes were made of borosilicate glass tubing (VWR micropipettes; VWR Co., West Chester, PA, USA) with DC resistance of 3-5 mΩ. Giga-Ohm seal resistance achieved after cell membrane break-in. The action potentials (APs) are recorded from differentiated NRVMs. The APs and APD were recorded under the current clamp configuration by Axon 700B amplifier (Axon Instruments, Irvine, CA, USA).

For APD recordings, the internal pipette solution contained 120 mM KCl, 10 mM EGTA, 10 mM Hepes, and 3 mM Mg-ATP, 1 mM MgCl_2_, at pH was adjusted to 7.2. The dose of 500 mg/mL Amphotericin B (Sigma) was included in the pipette solution. The external buffer contained 140 mM NaCl, 5.5 mM glucose, 5.4 mM KCl, 5 mM Hepes, 1.3 mM CaCl_2_, 0.5 mM MgCl_2_at pH 7.4 with NaOH. For whole Cell currents recording, the pipette and external solution were the same as APD’s.

For Ito1 recordings, the external solutions added with 200 mM CdCl_2_ to block Ca^2+^ currents. In order to avoid Na^+^-current contamination,the membrane potential was held at -40 mV or the NaCl in external solutions was substituted by equimolar choline [24]. Ito1 current was elicited from a holding potential of -40 mV to +100 mV in 10 mV increments by 300-ms depolarizing steps.

To test compound influence on native IKs currents, the Na^+^ in external solution was replaced byequimolar choline and 1 mM glibenclamide to suppress potential interference of IK1, INa, Ito1, ATP-dependent K+ channels (K-ATP), IKr and ICaL, respectively. The solution also was supplemented by 5 mM 4-AP, 1 mM dofetilide, 0.5 mM BaCl_2_, 0.2 mM CdCl_2_ [25]. IKs current was defined as the 10 mM chromanol 293B-sensitive current and elicited from a holding potential of -50 mV to potentials ranging from -50 mV to +100 mV in 10-mV increments by 3-s depolarizing steps.

For inward rectifier potassium currents (IK1) recording, the Na^+^ in external solution was replaced by equimolar choline and the solution was supplemented by 10 mM chromanol 293B-sensitive, 5 mM 4-AP, 1 mM dofetilide and 1 mM glibenclamide, 0.2 mM CdCl_2_ to minimize potential interference of INa, ICa, Ito1, IKr, IKs and ATP-dependent K^+^ channels (KATP), respectively [14]. Cells were held at -40 mV, and the currents were elicited by a cluster of potential from -120 mV to 0 mV (400 ms) in 10 mV increment.

For Cs^+^-carried IKr recording, the pipette solution contained 135 mM CsCl, 10 mM HEPES, 10 mM EGTA,and 5 mM ATP-Mg. The pH was adjusted to 7.2 with CsOH. The bath solution contained 135 mM CsCl, 10 mM glucose, 10 mM HEPESand 1 mM MgCl_2_. 10 mm nifedipine was added to delete potential interference of ICaL [26]. Cells were held at -80 mV, depolarizations in 10-mV increments to voltages between -70 and +70 mV for 1.5 s were applied to elicite currents. INa recording was conducted at room temperature, in a low-sodium extracellular solution containing 117.5 mM CsCl, 20 mM NaCl, 20 mM HEPES, 11 mM glucose, 1 mM CaCl2, 1 mM MgCl_2_, 0.1 mM CdCl2. The pipette solution contained 135 mM CsF, 10 mM EGTA,5 mM NaCl, 5 mM HEPES,5 mM Mg-ATP[27]. To characterize the voltage dependence of the peak INa, single cell was held at -120 mV, and 50 ms voltage steps were applied from -100 to +40 mV in 10 mV increments. Interval between voltage steps was 3 s.

For L-type calcium current (ICaL) studies, the external solution contained 136 mM TEA-Cl (tetraethylammonium chloride), 10 mM dextrose,10 mM HEPES, 5.4 mM CsCl, 2 mM CaCl_2_ and 0.8 mM MgCl_2_ (pH 7.4 with CsOH). The pipette solution contained 110 mM Cs-aspartate, 20 mM CsCl, 10 mM HEPES,10 mM EGTA, 5 mM Mg-ATP,1 mM MgCl_2_, and 0.1 mM GTP (pH 7.2 with CsOH) [28]. The ICaL currentwas determinedrepetitively at a test potential of 0 mV for 150 ms from Cells were held at -40 mV, and the currents were elicited by a cluster of potential from -50 to +50 mV in 5 mV increments.

### 2.5. Data analysis for Patch Clamp

All experiments data was recorded and analyzed by Clampfit 10.0 (Molecular Devices, Sunnyvale, CA, USA). Conductance–voltage (G-V) relationships was determined from peak current (I) versus different voltage by the Boltzmann equation: y= 1/(1 + exp[(V1/2 − V)/k]) in which V1/2, V, and k represented midpoint voltage of kinetics, test potential and slope factor, respectively. Data were analyzed by using GraphPad Prism 5. All data points are shown as mean ± S.E. n represents the number of the separate experimental cells.

## 3. Results

### 3.1. Toxicities of A. nagpag on Kunming mouse

Spiders inject venom to prey to quickly interfere with their physiology or kill them. We examined the toxic effects of venoms from *A. nagpag* against Kunming mouse. After intraperitoneal injection of 1 mg/kg venoms, mice exhibited classical symptoms of poisoning, such as the reduction of activity, posterior paresis and rapid breathing. At last, one mouse dead about 1 hour. At the concentration of 2.5 mg/kg or 5 mg/kg, the number of death mice are 2 and 4, respectively. When the concentration increased to 10 mg/kg, all the mice dead within 30 minutes. The LD_50_ value of *A. nagpag* on Kunming mice was 3.38 mg/kg (Table 1).

**Table 1.** Toxicities of *Alopecosa nagpag* on Kunming mouse.

| Concentration (mg/kg) | Number | Death | Mortality (%) |
| --- | --- | --- | --- |
| 0.5 | 6 | 0 | 0 |
| 1 | 6 | 1 | 16.67 |
| 2.5 | 6 | 2 | 33.33 |
| 5 | 6 | 4 | 66.67 |
| 10 | 6 | 6 | 100 |

### 3.2. The venom prolongs APD on isolated NRVMs

Venoms of *A. nagpag* showed mild toxicity to mouse. To further investigate toxicity, we examined effects of venoms on action potential duration (APD). APD of cardiomyocyte are divided into five phase including phase 0 to 4. At the dose of 100 μg/mL venoms of *A. nagpag* prolong APDs in phase 2 and 3 with a frequency-dependent manner. APDs were significantly altered at the frequency of 1 Hz, while only slightly changes were detected at 2Hz (Fig. 1).

**Figure 1.**
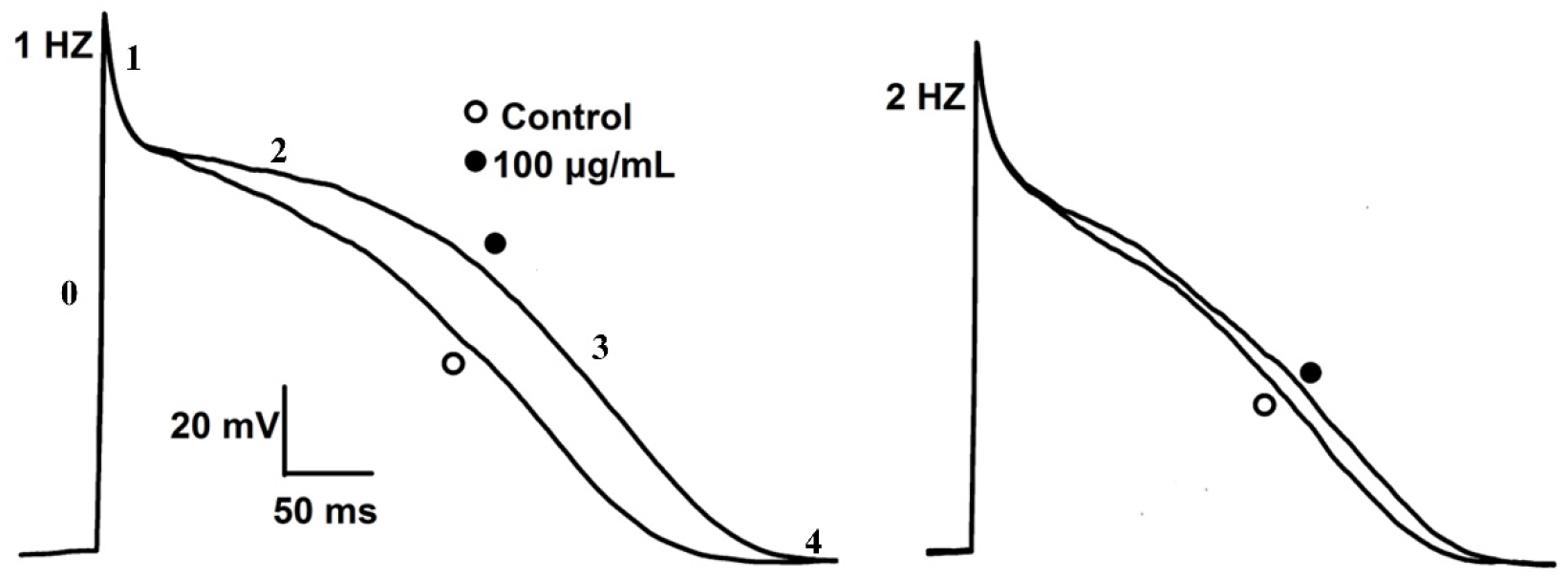
Effects of the venom of *A. nagpag* on action potentials in NRVMs. The action potentials elicited at 1 (left) or 2Hz (right) in thecontroland 100 μg/mL venom.

### 3.3. Effects of venoms on whole currents of NRVMs

Membrane currents were evoked using a short voltage ramp from −80 mV to +50 mV at a holding potential of −80 mV (Fig. 2, insert). Control current trace was consistent with previous work [7]. All ion currents were functionally expressed in the cardiac cell, including I_K1_, I_Ks_, I_to1_, I_Kr_, I_CaL_ and I_Na_. Compared with control currents, phase II of whole ion currents was inhibited by 100 μg/mL venoms (Fig. 2). Phase III of whole ion currents was significantly enlarged after treatment with venoms. According to the results, we proposed that I_k1_, I_Na_ and I_CaL_ currents were altered by 100 μg/mL venoms. Effects of venoms on typical cardiac ion currents (I_K1_, I_Ks_, I_to1_, I_Kr_, I_CaL_ and I_Na_) were further studied.

**Figure 2.**
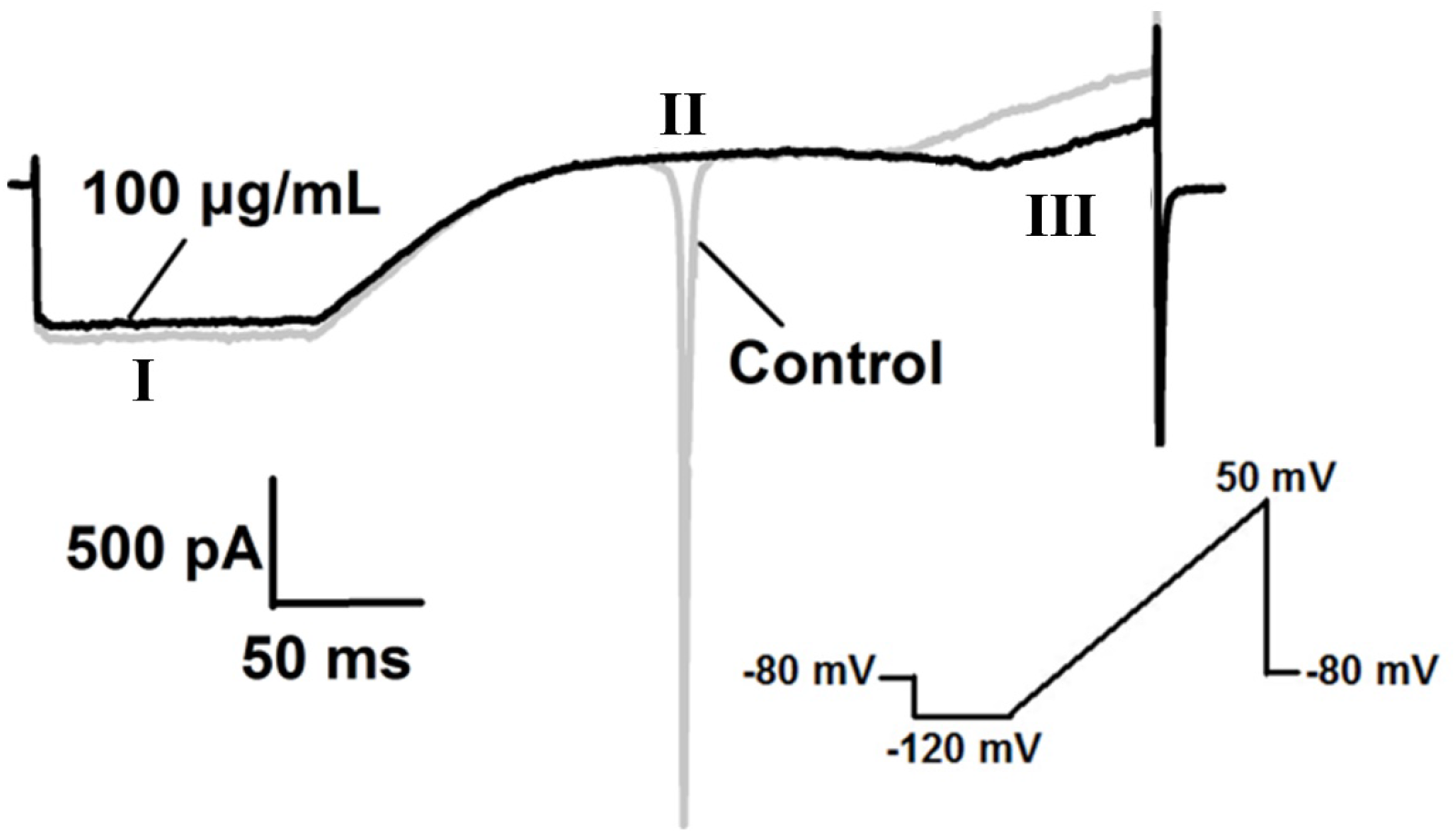
Effects of the venom on whole cell currents of NRVMs. The holding potential was −80 mV, and the shape of the voltage ramp is shown at the below. Gray line shows the control, black line shows the experimental group of 100 μg/mL of venom.

### 3.4. The venoms inhibit INa on NRVMs

Sodium channel plays an important role in cardiac conduction and generates the fast depolarization of cardiac action potential[8]. Sodium channel currents of rat ventricular myocytes were elicited from a holding potential of -80 mV by a 50 ms depolarizing to -10 mV. Currents were almost completely inhibited by 100 μg/ml venoms (Fig. 3C). The current-voltage (I-V) of rat ventricular myocytes was activated about -40 mV and revised about +20 mV. No shift was detected on the I-V curve before and after 100 μg/ml *A. nagpag* venom treatment (Fig. 3A, B, D), indicating that the toxicity was not associated with the I-V relationship of the cardiac I_Na_.

**Figure 3.**
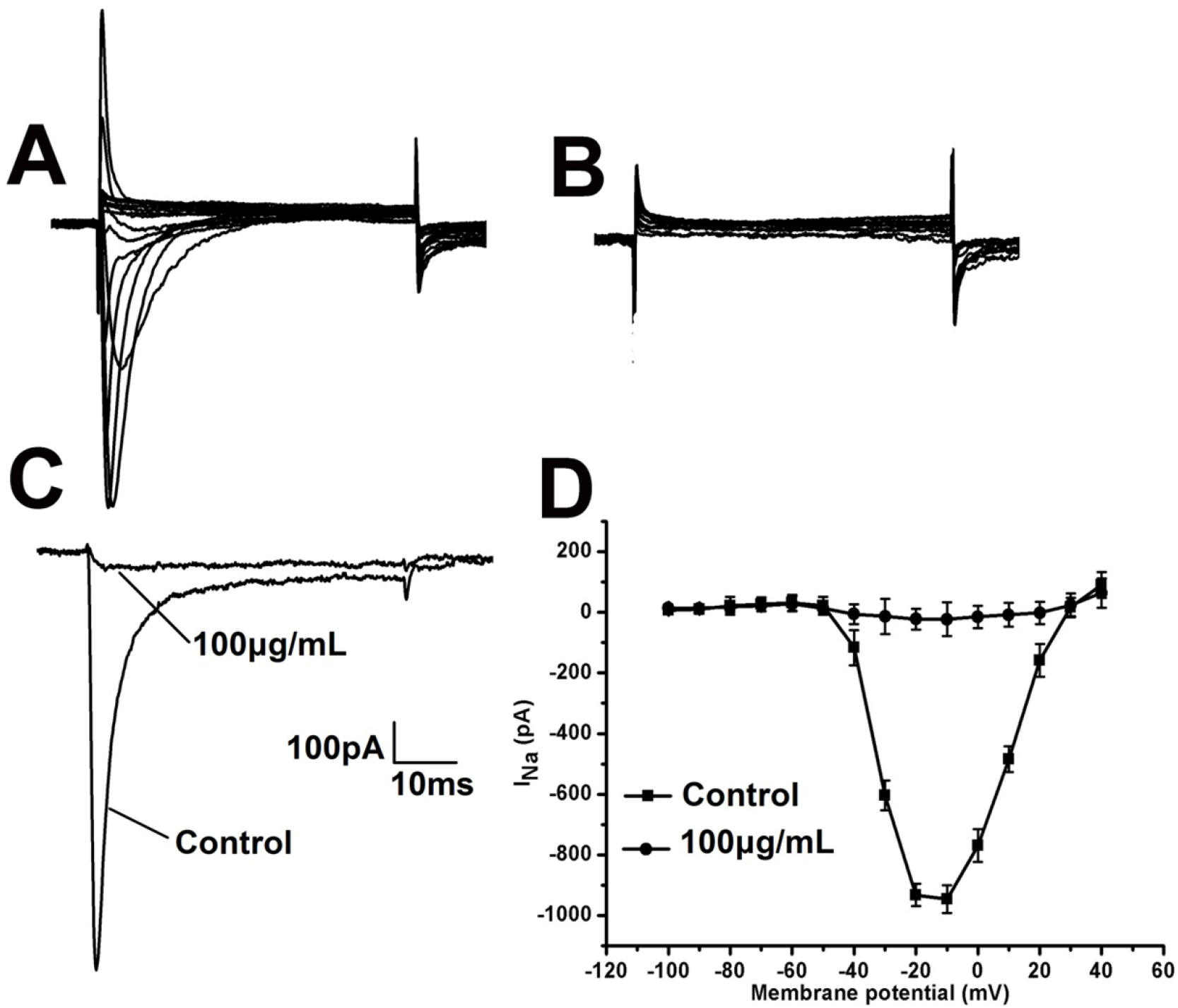
Effects of the venom on I_Na_ recorded in NRVMs. Currents were elicited by voltage steps from a holding potential of -80 mV. A and B, representative recording of currents in the absence and presence of 100 μg/mL of venom t. C, The application of 100 μg/mL of venom inhibited I_Na_ currents. D, Effects of 100 μg/mL of venom on average steady-state current-voltage (I-V) relationship.

### 3.5. Activity of the venoms on ICaL

We elicited I_CaL_ using a test potential of 0 mV with a holding potential of -40 mV. After the first I_CaL_ trace elicited, 100 μg/mL venoms were added to the cell to observe the change of I_CaL_ currents. Venoms largely increased I_CaL_ by 132 ± 4.0% (Fig. 4C). Typical I-V traces of calcium currents on rat ventricular myocytes were shown in Fig. 4 A and B. The I-V curve did not evidently alter after treating with100 μg/mL venoms (Fig. 4D).

**Figure 4.**
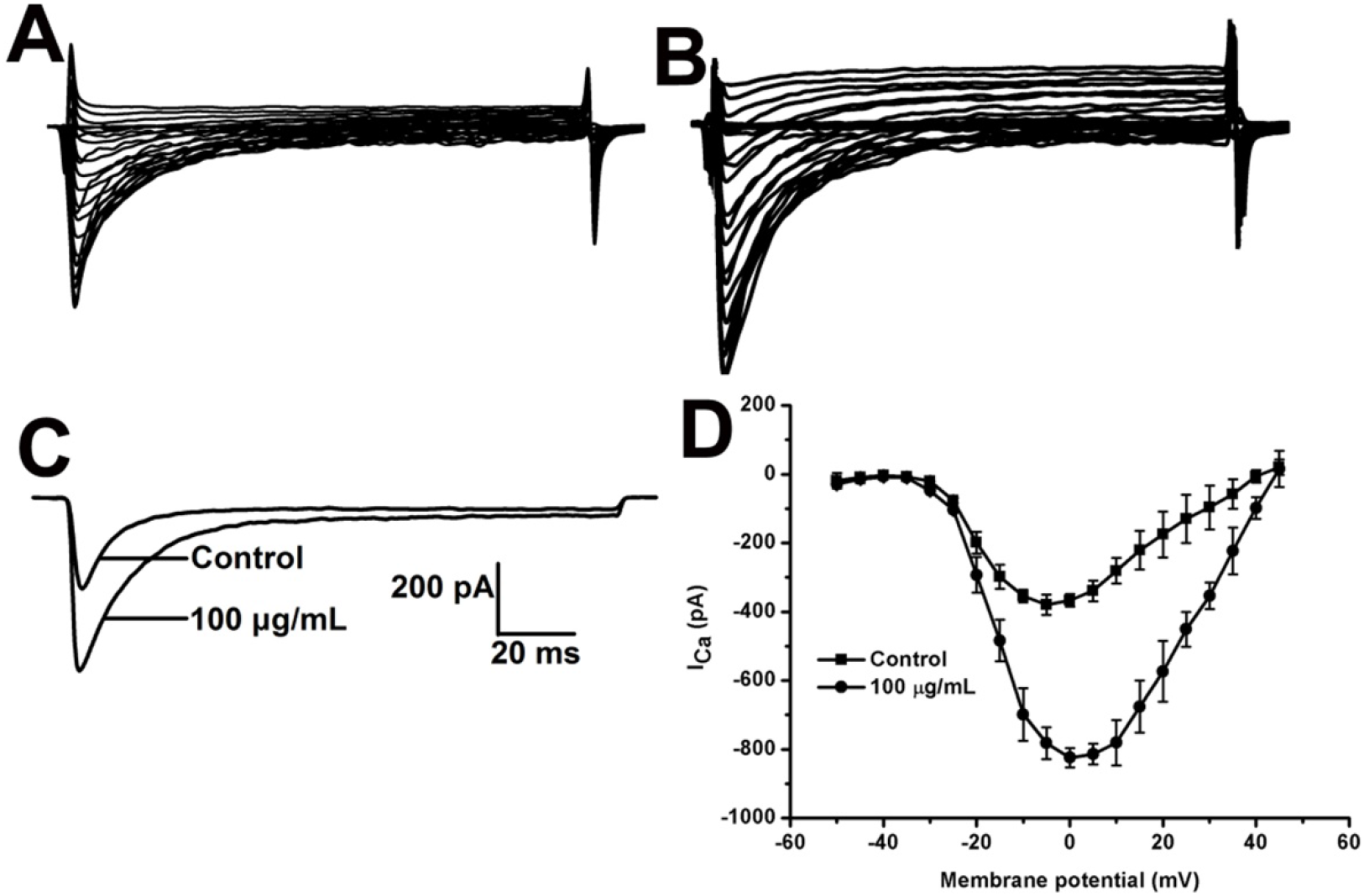
Effects of the venom on I_CaL_ recorded in NRVMs. Currents were elicited by voltage steps from a holding potential of -40 mV. A and B, representative current traces recorded in the absence and presence of the venom (100 μg/mL). C, 100 μg/mL venom enhanced I_CaL_. D, Effects of 100 μg/mL of venom on average steady state current-voltage (I-V) relationship.

### 3.6. Effects of the venoms on ventricular repolarizing currents IKs, Ito1, IK1 and IKr

There are two distinct current components of I_K_ on rat ventricular myocytes, including slowly activating delayed rectifier outward K^+^ currents (I_Ks_) and rapidly activating delayed rectifier outward K^+^ currents (I_Kr_). To independently record I_Ks_, a class III antiarrhythmic agent and a selective blocker dofetilide of I_Kr_ were used to inhibit I_Kr_ currents [9]. I_Ks_ currents of NRVMs were evoked by a 3-s-long voltage-clamp pulse protocol. 100 μg/mL venoms did not produce significant inhibition in I_Ks_ currents (Fig. 5C). Meanwhile, venom did not alter the I-V curves (Fig. 5D).

**Figure 5.**
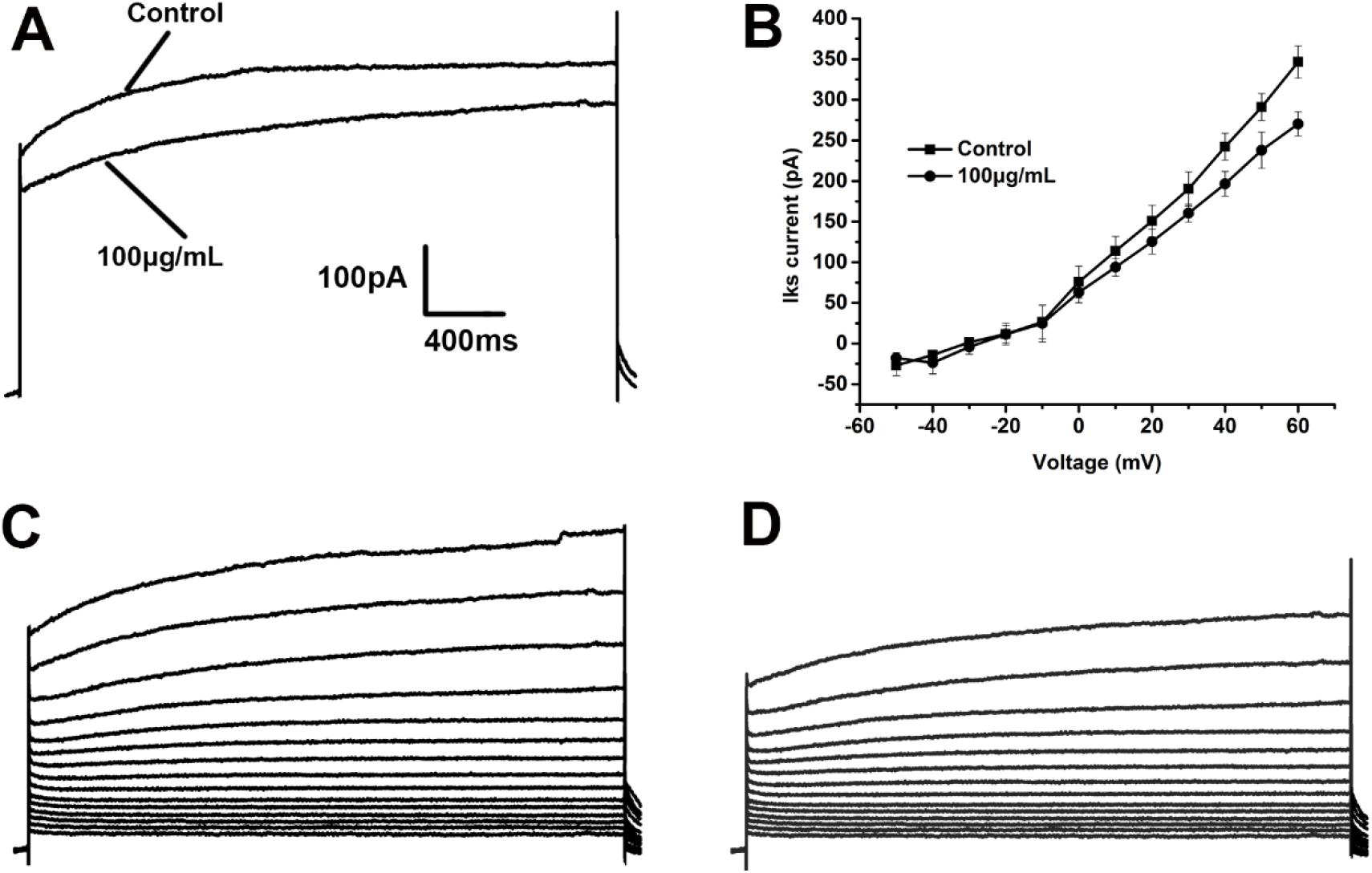
Influences of the venom on I_Ks_ recorded in NRVMs. Currents were elicited by voltage steps from a holding potential of -40 mV. A, 100 μg/mL venom inhibited I_Ks_. B, effects of the venom on normalized steady-state current-voltage (I-V) relationship. C and D, representative recording of currents in the control and 100 μg/mL venom ().

The transient outward potassium current (I_to1_) plays a key role in action potential (AP) shape and dynamics. I_to1_ also contributes to cardiac arrhythmia by decreasing the AP plateau voltage to a low range at which I_CaL_ is proper for reactivation and I_Ks_ is not available to activate [10]. At the concentration of 100 μg/mL, *A. nagpag* venom showed a little effect on I_to1_ with current amplitude increasing 9.6 ± 3.6% (Fig. 6C). The activation of I_to1_ before and after adding venom also exhibited no significant changes in I-V curve. The half-activation voltage in the presence of the venom was 17.6 ± 1.5 mV, compared to 23.6 ± 1.4 mV in the control (Fig. 6E, n=5).

**Figure 6.**
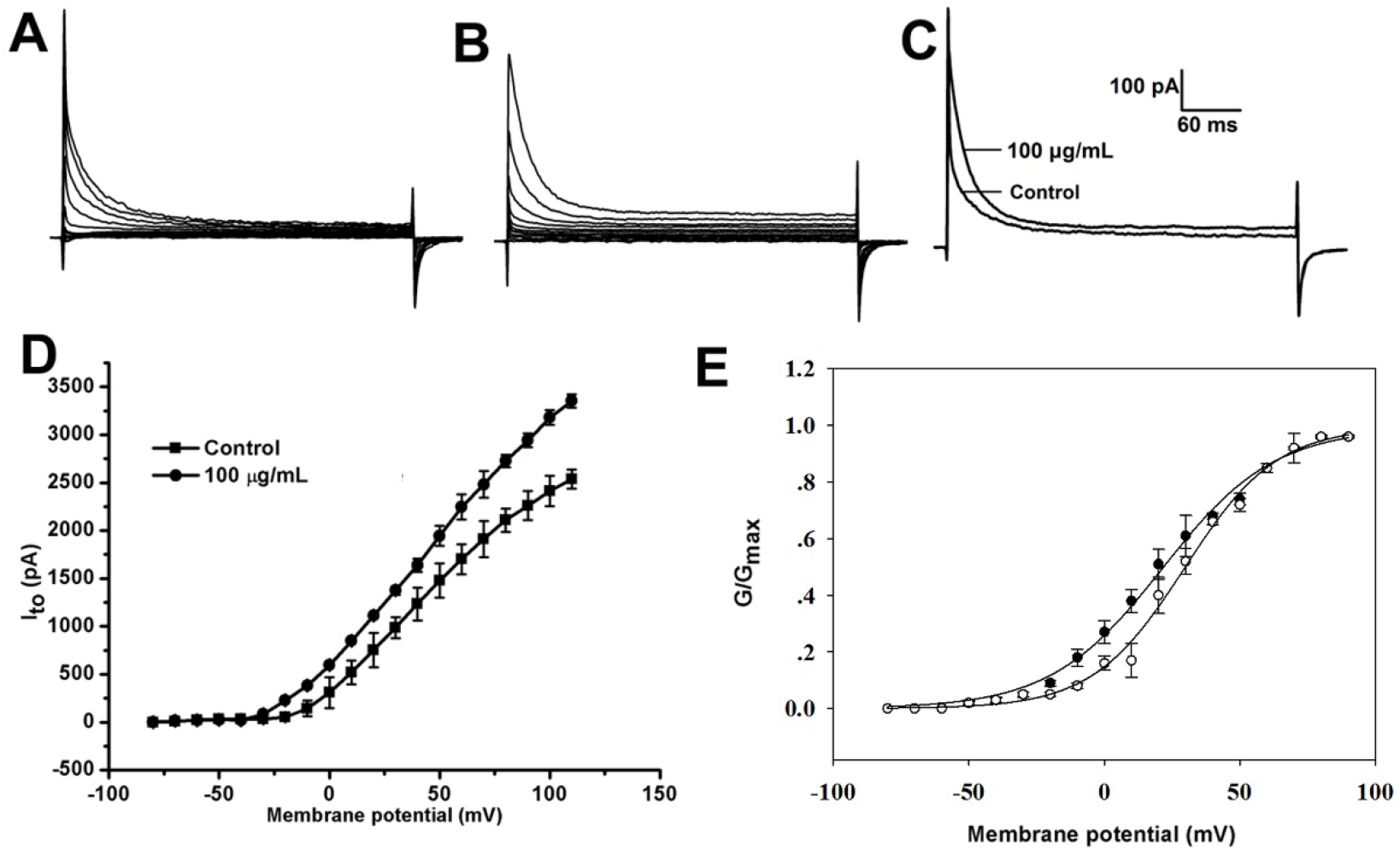
Effects of the venom on I_to_ recorded in NRVMs. Currents were elicited by voltage steps from a holding potential of -40 mV. A and B, representative recording of currents in the control and 100 μg/mL venom. C, effects of100 μg/mL venom on I_to1_. D and E, effects of the venom on normalized steady-state current-voltage (I-V) relationship and G-V relationship.

Inward rectifier potassium current (I_K1_) participates in the late phase repolarization of ventricular action potential and is an important element relating to resting membrane potential [11]. Abnormality of I_K1_ may result in heart failure and arrhythmias. As shown Fig. 7C, little inhibition could be found in the traces before and after 100 μg/mL venoms treatment. The venoms only activated I_K1_ in NRVMs by 12.4 ± 3.7% (n > 6) and with no obvious change in the current-voltage relationship (Fig. 7C, D).

**Figure 7.**
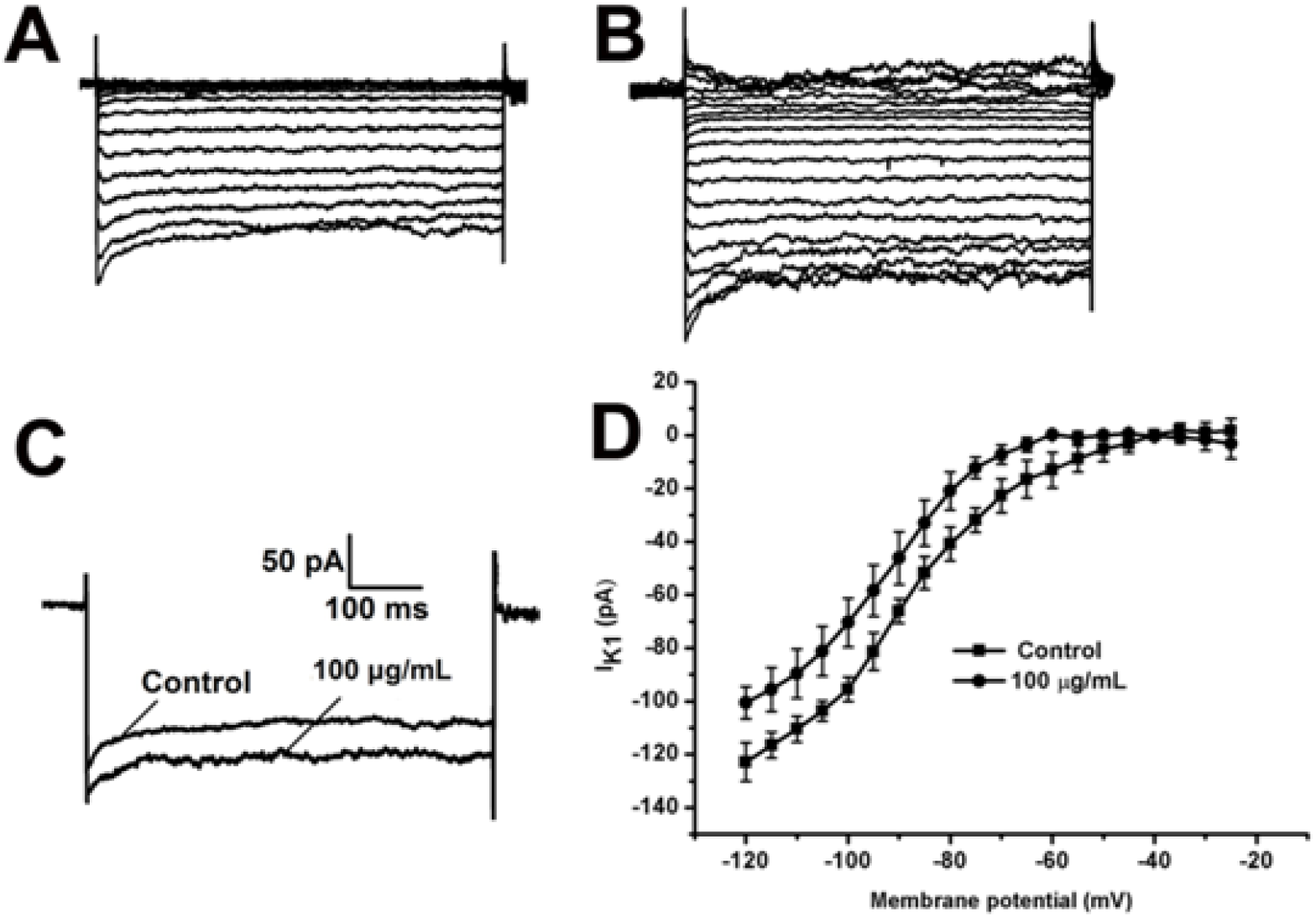
Effects of the venom on I_K1_ recorded in NRVMs. Currents were elicited by application of voltage steps from a holding potential of - 40 mV. A and B, representative trace recordings of currents in the control and 100 μg/mL venom. C, 100 μg/mL venom decreased I_K1_. D, Effects of the venom on normalizedsteady-state current-voltage (I-V) relationship.

Rapidly activating delayed rectifier potassium current (I_Kr_), namely as hERG currents, is vital for cardiac repolarization. Reduction of I_Kr_ may cause long QT syndrome (LQTs) and prolong APD, finally resulting in fatal arrhythmias or sudden death [12]. Recent study showed that I_Kr_ channels display unique Cs^+^ permeability [13]. Therefore, we use isotonic Cs^+^ solution to record pure I_Kr_ in NRVMs as described previously [14]. Cs^+^-carried I_Kr_ currents were elicited by a 10-mV depolarization from a holding potential of -80 mV. The following tail currents at -80 mV displayed an initial rising phase, indicating a rapid recovery of inactivated channels to the open state before deactivation, and is unique to I_Kr_. As shown in Fig.8, 100 μg/mL venom obviously decreased the peak and tail I_Kr_ current of ventricular myocytes. The inhibition of peak currents and the tail currents by venoms were 54.3 ± 3.2% and 60.8 ± 4.3%, respectively (Fig. 8C). The I-V relationships of peak currents and the tail current activation curves showed no obvious change (Fig. 8D, E).

**Figure 8.**
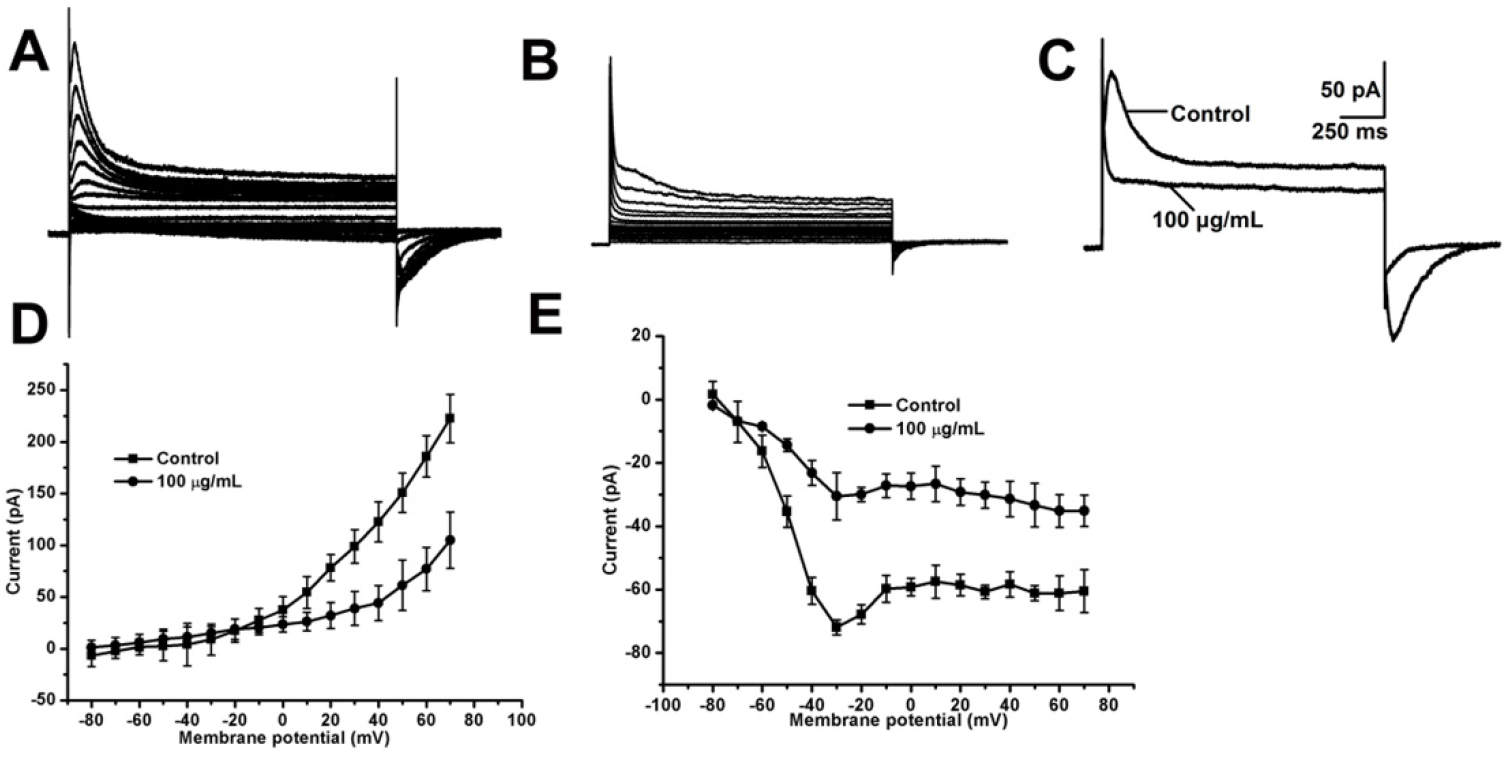
Cs^+^ currents recorded in NRVMs with both pipette and bath solutions containing 135 mM Cs+. A and B, Representative recording of currents in the control and 100 μg/mL venom 100 μg/mL. C, By depolarization to voltages +50 mV from the holding potential -80 mV, the Cs+ currents were elicited in the controland 100 μg/mL venom. D, influence of the venom on normalized steady-state current-voltage (I-V) relationship of the maximal current during depolarization. E, effects of the venom on normalizedI-Vrelationship of the current at the end of depolarizing steps.

## 4. Discussion

Most of spiders use venoms to immobilize or lead function disorder of their preys. Some spiders show high toxicity to mouse, for example LD_50_ value of *Ornithoctonus hainana* (*O. hainana*) on Kunming mouse is only 1.25 mg/kg. Venoms of *O. hainana* could prolong APD of cardiac myocytes [15]. It implies that the toxicity effects of *O. hainana on* cardiac myocytes may be one reason why spider could cause mouse to death. Besides *O. hainana*, toxins from *Chilobrachys jingzhao (C. jingzhao)* also could act on ion channel of cardiac myocytes. JZTX-II and JZTX-III, both isolated from the venom of *Chilobrachys jingzhao* (*C. jingzha*o), inhibit sodium currents of cardiac myocytes (Nav1.5) with high affinity [16]. Venoms of *A. nagpag* showed mild toxicity to Kunming mouse with LD_50_ of 3.32 mg/kg, implying venom could significantly disorder the physiological function of prey. Then, we found that APD of NRVMs was prolonged in a frequency-dependent manner. Meanwhile, whole ion currents of NRVMs were also inhibited by 100 μg/mL venom. Thus, venoms of *A. nagpag* should contain toxins acting on ion channel.

Phases of the cardiac action potential differ from the neuronal action potential and divide into five phases (phase 0-4). If the membrane potential of cardiac is raised up to the threshold, fast Na^+^ channels are activated and initiated upstroke of the AP. The sharp rise in voltage (phase 0) corresponds to the influx of sodium ions. To our surprise, 100 μg/mL venom inhibited I_Na_ currents, but no obvious changes were found in Phase 0. A brief rapid repolarization makes cells enter to phase 1 due to potassium ion though rapid activating and inactivating transient outward K^+^ channel (I_to_). Venoms showed a little effect on I_to_, so no changes were detected on phase 1. The extended plateau (phase 2) is a distinguishing feature of cardiac action potential, which results from opening of voltage-sensitive calcium channels for a few hundred milliseconds [17]. Venoms could increase Ca^2+^ currents and phase 2 was significantly prolonged due to more Ca^2+^ ions to cardiac ventricular myocytes though L-type calcium channels. Activation of delayed-rectifier currents, particularly the rapid delayed-rectifier I_Kr_, determinate the action potential with an appropriate delay by producing rapid phase 3 repolarization. Phase 3 was also prolonged because I_Kr_ are significantly decreased by 100 μg/mL venoms. Venoms prolong APD phase 2 and 3 may lead a sudden chaos of cardiac rhythm and cause preys to death, because the cardiac action has a direct relation with the heart cardiac rhythm. Our previous work has demonstrated that *O. hainana* venom also could change the cardiac rhythm of Kunming mouse and exhibits a LD_50_ of 1.25 mg/kg on Kunming mouse [18]. Thus, change of cardiac rhythm may be one strategy that spiders use to in order to kill prey.

*A. nagpag* is a small-size wolf spider disturbed in Yunnan province of China, which preys mainly on insects and sometimes on mammals. Small-size wolf spiders have long been known to be venomous, but their venoms have been little studied. In this study, we demonstrate *A. nagpag* shows effect on cardiac ion currents of ventricular myocytes and could kill mouse. It implies that the venoms should contain many cardiac channel antagonists and might be a rich source for drug development. The current study will stimulate more interest in the venoms of this hitherto neglected group, small-size wolf spider.

## Ethical statement

All the experiments were reviewed and approved by the Committee on the Use and Care of Animals at the Yunnan Province, People’s Republic of China and were conducted in accordance with the guidelines established by the Committee.

## Acknowledgments

We are thankful to Dr. mingqiang Rong for knowledge support. This work was supported by the Chengdu Pepbiomedical Co., Ltd.and Animal Center of Kunming Institute of Zoology.

## AUTHORS’ CONTRIBUTIONS

Z.X.G. conducted experiments on sample preparation and performed most electrophysiological. Z.X.G., C.B.L., YHL., and B.H. designed the experiments, analyzed the data, and wrote the manuscript. All authors read and approved the final version of the manuscript.

